# Serial dependence in timing at the perceptual level being modulated by working memory

**DOI:** 10.1101/2022.11.11.516083

**Authors:** Shuai Chen, Tianhe Wang, Yan Bao

**Author notes:** Correspondence: Dr. Yan Bao, School of Psychological and Cognitive Sciences, Peking University, 5 Yiheyuan Road, Beijing 100871,China.

## Abstract

Recent experiences bias the perception of following stimuli, as has been verified in various kinds of experiments in visual perception. This phenomenon, known as serial dependence, may reflect mechanisms to maintain perceptual stability. In the current study, we examined several key properties of serial dependence in temporal perception. Firstly, we examined the source of serial dependence effect in temporal perception. We found that perception without motor reproduction is sufficient to induce the sequential effect; the motor reproduction caused a stronger effect and it is achieved by biasing the perception of the future target duration rather than directly influences the subsequent movement. Secondly, we ask how the working memory influences serial dependence in the in a temporal reproduction task. By varying the delay time between standard duration and the reproduction, we showed that the strength of serial dependence enhanced as the delay increased. Those features of serial dependence are consistent with what has been observed in the visual perceptual tasks, for example, orientation perception or location perception. The similarities between the visual and the timing tasks may suggest a similar neural coding mechanism of magnitude between the visual stimuli and the duration.

## Introduction

Our perception of the external world is remarkably stable over time despite the rapid changes of physical attributes of sensory stimuli. Past experience influences the perception of current stimuli. This sequential dependence produces an auto-correlation creating a stream of continuous perception. The temporal stability is presumably based on a mechanism that is experimentally reflected in serial dependence of stimulus processing (Fischer & Whitney, 2014). Serial dependence has been demonstrated in different studies using various visual stimuli, e.g., spatial position (Papadimitriou et al., 2015), motion direction (Alais et al., 2017), numerosity (Pascucci et al., 2019), direction of eye gaze (Alais et al., 2018), and some of the higher level attributes like facial expression (Liberman et al., 2014) or attractiveness(Xia et al., 2016), and even in complex judgements such as variance (Suárez-Pinilla et al., 2018) and confidence (Rahnev et al., 2015). All these results point out that serial dependence my serve as a general principle on different levels at least in visual perception.

Similar serial dependence effect has also been observed in timing (Wang et al., 2021). Our previous study showed that current reproduced duration is biased towards the one back standard duration in the temporal reproduction task. Similar to the effect in the visual perceptions, this attractive bias showed a non-linear relationship with the difference between the current and the previous standard durations: the bias increases with the difference when it is relatively small and then decreases when the difference between durations is too large.

However, when we tried by examine the source of this sequential effect, serial dependence is almost neglectable if participants made perception without performing a reproduction. Similar to this result, other studies in temporal perception also showed that the effect of immediately-prior experience will extinguish if the prior interval was not linked to a motor command (Cai et al., 2017; Dyjas et al., 2012; Wehrman et al., 2018). Given these results, it is unclear whether serial dependence in timing actually rests in perception or it could be a pure motor effect, that is, the current motor reproduction biases the subsequent reproduction without changing the perception of the standard duration. It should be noted that in the previous study evaluating the relative contribution of perception and motor reproduction (Wang et al., 202), participants were exposed to the duration twice in the action trial (perception and reproduction) but only once (perception) in the no-action trial. This difference might result in a weakened effect in no-action trials, thus serial dependence might not have been captured in those trials.

To reexamine the question that whether perception without movement can induce serial dependence, in experiment 1, we introduced a temporal reproduction task where we repeated non-action trials for 1-5 times between each two action trials. Moreover, in a continuous temporal reproduction task, participants are required to make motor responses on each trial, thus it is not sufficient to answer the question whether it is the motor response or whether the perception of the current trial is influenced by the prior trial. To this end in experiment 2, we combined a temporal reproduction task with a temporal comparison task, the rationale being that by using a temporal comparison task to probe serial dependence, we rule out the possibility of the effect of motor responses between subsequent trials. The preceding reproduction task provides a temporal context for the following comparison task. If the serial dependence induced by the reproduction task works as changing the perceptual representation of the current stimuli, then we would observe that the subjective equivalence point will be biased by the preceding reproduction task.

Besides the source of serial dependence, in experiment 3, we examine whether the working memory influence serial dependence. In visual perception, a number of studies showed that increasing the interval between perception and reproduction enhances the serial dependence effect, perhaps because the uncertainty to the current perception increases across time so that during retrieval the neural system rely more on the prior information. It is interesting to ask whether serial dependence in duration perception is modulated in a similar way. If serial dependence increases over longer delays in temporal tasks, it might suggest a similar mechanism for duration and visual stimulus processing during their retention period in working memory; this would further validate the argument that the serial dependence effect is the result of a generalized processing mechanism.

In summary, there were three primary goals of the current studies: In experiment 1 we aimed to disentangle different cognitive factors that could potentially influence serial dependence. In experiment 2 we developed a novel two-stage paradigm to examine whether serial dependence could be transferred between different component, e.g., from reproduction to perception. In experiment 3 we sought to assess the effect of a delay period on the serial dependence in a temporal reproduction task.

### Experiment 1

Previous research has shown that in temporal reproduction tasks, the serial dependence bias (Wang et al., 2021) was mainly contributed by the reproduction other than the perception. However, it should be noted that in the prior experiment examining the serial dependence effect, participants were exposed to the duration twice in the action trial (perception and reproduction), but only once (perception) in the no-action trial. This difference might cause different results in the analyses of action and no-action trials. Moreover, serial dependence originating from perception was possibly not captured in the prior experiment perhaps due to the insensitive measurement used. Thus, to amplify the potential serial dependence effect caused by perception, here we increased the percentage of no-action trials.

## Materials and Methods

The first experiment was designed to disentangle the motor and perceptual component in serial dependence.

### Participants

Fourteen healthy volunteers (8 males and 6 females, 18-30 years old, mean age = 21.7 ± 4.00) with normal or corrected to normal vision participated in Experiment 1. All subjects reported not taking any psychotropic drugs before the experiment. Before the experiment start, all participants had given written informed consent; the study was approved by the institutional review board at Peking University. Subjects were screened for red-green color blindness before conducting the experiment. Financial compensation was given upon the completion of the experiment.

### Apparatus

The visual stimuli were presented on a 27-inch monitor equipped with a GeForce 1080 graphics card. The computer was equipped with Windows 8 operating system, with a refresh rate of 100 Hz, and a resolution of 1024 × 768 pixels. Participants were seated 65 cm from the monitor in a dark soundproof room, participants’ heads were stabilized using a chin rest. The brightness of the screen was controlled in an appropriate range and kept constant, and a Dell keyboard is used for response.

### Stimuli and procedure

Each trial started with a fixation point at the center of the screen. After a random interval ranging from 0.7-1.2 s (following an exponential distribution), a ripple-shaped stimulus centering at the fixation point (see Fig.1A) was presented for sample duration and then disappeared. After 300ms, the fixation point changed into either a “+”, that informed participant to reproduce the duration of the stimulus (action trial), or a “x”, which informed participants to make no response (no-action trial). In the action trials, participants could make a response at any time they want by holding the keypress for sample duration. After the key released, the fixation point changed color to present feedback for 300 ms. In the no-action trial, the “x” presented for 700 ms. Then the fixation point changed back into a point indicating the start of a new trial.

**Figure1.**
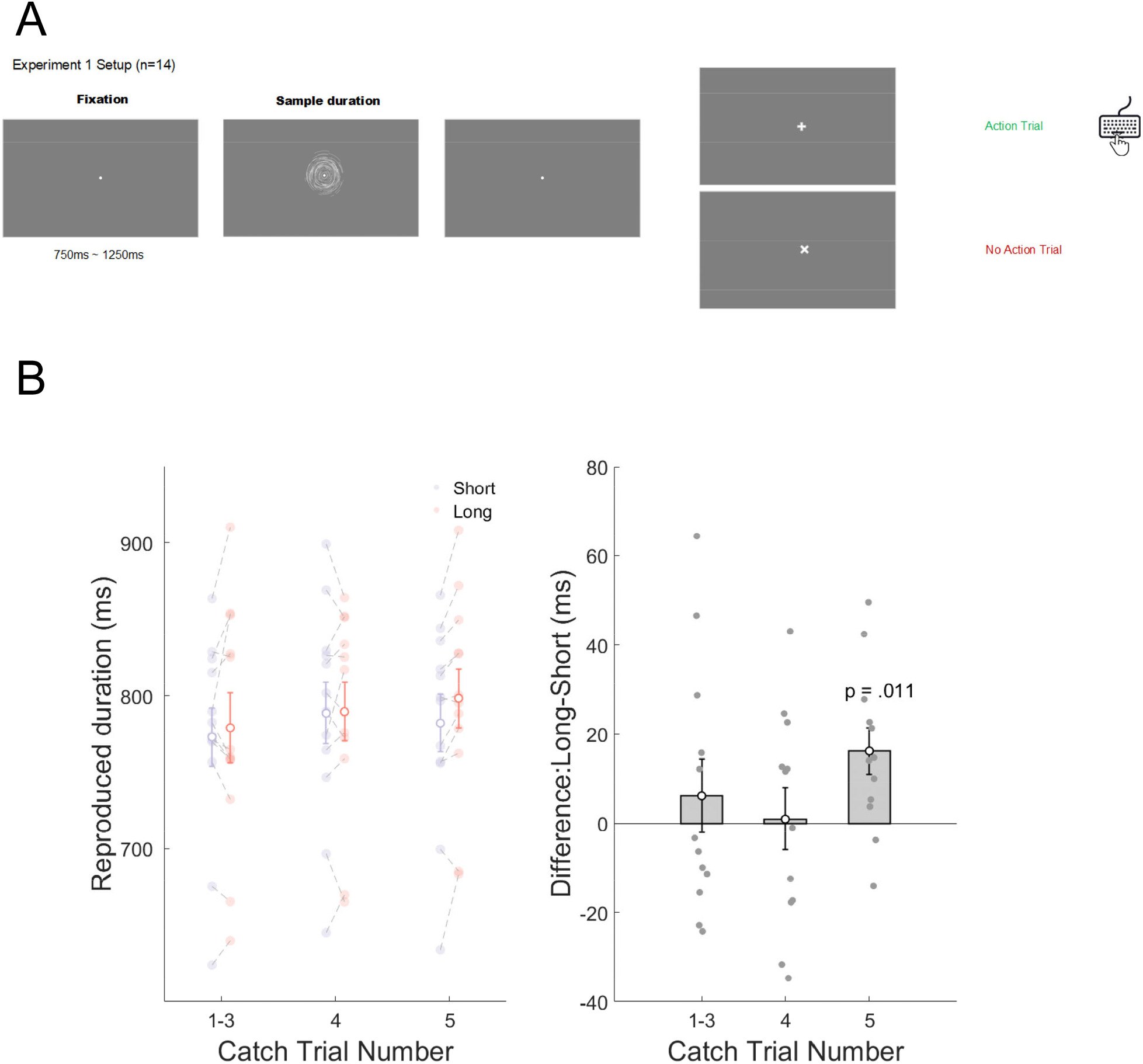
(A) Schematic procedure of experiment 2. A ripple-shaped stimulus was presented for sample duration. After it disappeared, the fixation point will change into one of following two shapes, “+” indicating that participants should hold a keypress to reproduce the stimulus (action trial) and “x” indicating not to make a response (no-action trial). Participants were instructed to press and hold the key for a period that matching the sample duration. After the key was released, the fixation point changed color to give feedback. In the no-action trial, a new trial started after presenting the “x” for 700 ms.(B) Result of experiment 2. The left panel shows the reproduction time under different preceding conditions (long or short). Each colored dot represents one subject. Error bar represents standard error. The right panel calculates the difference of reproduced durations between the long and short prior conditions across different numbers of no-action trials preceded the current action trial. Error bar represents standard error.

The sample duration of action trials was fixed at 700 ms. Zero to five no-action trials sampled from either the long or short prior conditions were inserted between every two action trials. The short prior condition contained 500, 550, 600, 650 ms. The long prior condition contained 750, 800, 850, 900 ms. All eight sample durations in short and long prior conditions appeared for equal times in the experiment. The probability of 0, 1, 2, 3, 4, 5 no-action trials between two action trials was 10%, 10%, 10%, 10%, 30%, 30%, respectively. Participants went through a total of 2064 trials, among which 480 trials were action trials. Prior conditions were pseudo-randomized within blocks of 516 trials (120 action trials), with a rest of one minute between every two blocks.

### Analysis

The reproduced durations shorter than 0.3 s or longer than 1.5s (less than 1‰) were seen as outliers and excluded from further analysis. Action trials were grouped based on the number of no-action trials ahead of them and whether those no-action trials belonged to the long prior or the short prior conditions. Action trials after 1, 2, and 3 no-action trials were grouped together to make each no-action trial number condition (1 to 3, 4, 5 no-action trials) contain 30% of action trials. The difference between the reproduced durations in long and short prior conditions was calculated for different numbers of no-action trials, respectively. One sample t-tests were applied to compare the differences from zero.

As for the calculation of serial dependence, we applied an index “deviation” as in previous research to quantify the strength of response error (Wang et al., 2021). The deviation was calculated by subtracting the mean reproduced duration of the corresponding standard interval from reproduced duration on each trial. The purpose was to detrend the influence of central tendency effect from all trials.

To quantify the strength of serial dependence, we fit the Derivate of Gaussian(DoG model; Fischer & Whitney, 2014) to our data:

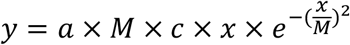

The instantaneous slope of the DoG curve indicates the direction of the sequential effect, with positive values implies an attractive effect and negative values implies a repulsive effect. To evaluate the strength of serial dependence, we used the peak-to-peak value (p2p) in the DoG curve as a measure.

## Results

We first divide the data according to the number of no-action trials ahead of them and the type of no-action trials (long prior condition or short prior condition). We confirmed that no participants realized that the sample duration in the action trial was fixed from the post-experiment inquiry. As shown in figure 1.B, when preceded by 1-4 no-action trials, no significant difference between reproduced durations was found between long and short prior conditions (1∼3 no-action trials: t_(11)_ = 0.76, p = 0.46; 4 no-action trials: t_(11)_ = 0.14, p = 0.89). However, the reproduced duration in the long prior condition was significantly longer than that in the short prior condition when preceded by five no-action trials (difference:16.14 ± 18.20 ms, t_(11)_ = 3.07, p = 0.011). It should be noted that though the effect is significant, its magnitude is much smaller compared to the serial dependence effect in experiment 1. These results suggested that the perceptions in the no-action trials weakly attracted the reproduction in the action trial, and it can only be observed when the effect has been accumulated, such as five no-action trials preceded before the current action trial.

### Experiment 2

It is possible that the attractive effect we observed in experiment 1 was due to motor hysteresis, i.e. individuals tend to reuse a recently used motor response rather than generating a new one. To answer this question, we need to include a non-motor task to capture this bias. Here we also want to test whether the bias from a reproduction is task specific and it can also influence a different subsequent task, e.g., a temporal comparison task. Thus, we developed a two-stage paradigm to answer these questions.

## Materials and Methods

### Participants

Sixteen healthy volunteers (6 males and 10 females, 18-30 years old, mean age = 22.87 ± 2.26) with normal or corrected to normal vision participated in Experiment 3. All subjects reported not taking any psychotropic drugs before the experiment. Before the experiment start, all participants had given written informed consent; the study was approved by the institutional review board at Peking University. Financial compensation was given upon the completion of the experiment.

### Apparatus

The visual stimuli were presented on a 27-inch monitor equipped with a GeForce 1080 graphics card. The computer was equipped with Windows 10 operating system, with a refresh rate of 100 Hz, and a resolution of 1920 × 1080 pixels. Participants were seated 70 cm from the monitor in a dark soundproof room, participants’ heads were stabilized using a chin rest. The brightness of the screen was controlled in an appropriate range and kept constant, and a Dell keyboard is used for response.

### Stimuli and procedure

The experimental procedures were generated using MATLABR2021b and PsychToolBox 3.0.14 toolkit (Mathworks, Natick, MA; Brainard 1997; Pelli 1997). A scheme of the task is shown in Figure 2.A. Each trial includes two phases. In the first phase of each trial, a gray center fixation point with 0.5 degrees in diameter was presented on a black screen. After a period of 0.5s to 1s, a ripple shape stimulus was presented on the screen for a certain standard duration. The subjects were instructed to reproduce the intervals with their dominant hands. The standard interval ranged from 400 ms to 1200 ms in five arithmetical equal steps. After the reproduction, the central fixation point changed into a cross, indicating the beginning of the second phase. After 500 ms delay, a ripple shape pattern with a standard duration of 800 ms appeared on the screen. We add 7% of catch trials, in which the standard duration was either 400ms or 1200ms. The purpose of catch trials was avoiding participants remembering the standard duration instead of attending to it. Those catch trials were discarded in further analysis. After another 500 ms, a ripple shape pattern with a comparison duration appeared. The comparison duration compromised of five durations ranging from 400 ms to 1200 ms in five arithmetical equal steps. The task in second phase was to indicate is the second duration longer or shorter than the first duration by pressing “s” and “l” key respectively. To promote for better performance, a feedback of block accuracy was presented after each block. The reproduction accuracy was calculated by 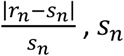 presents the standard duration of the *n*th trial and *r_n_* represents the reproduced duration of the *n*th trial. The comparison accuracy was indexed by the correct responses divided by the total trials.

**Figure 2.**
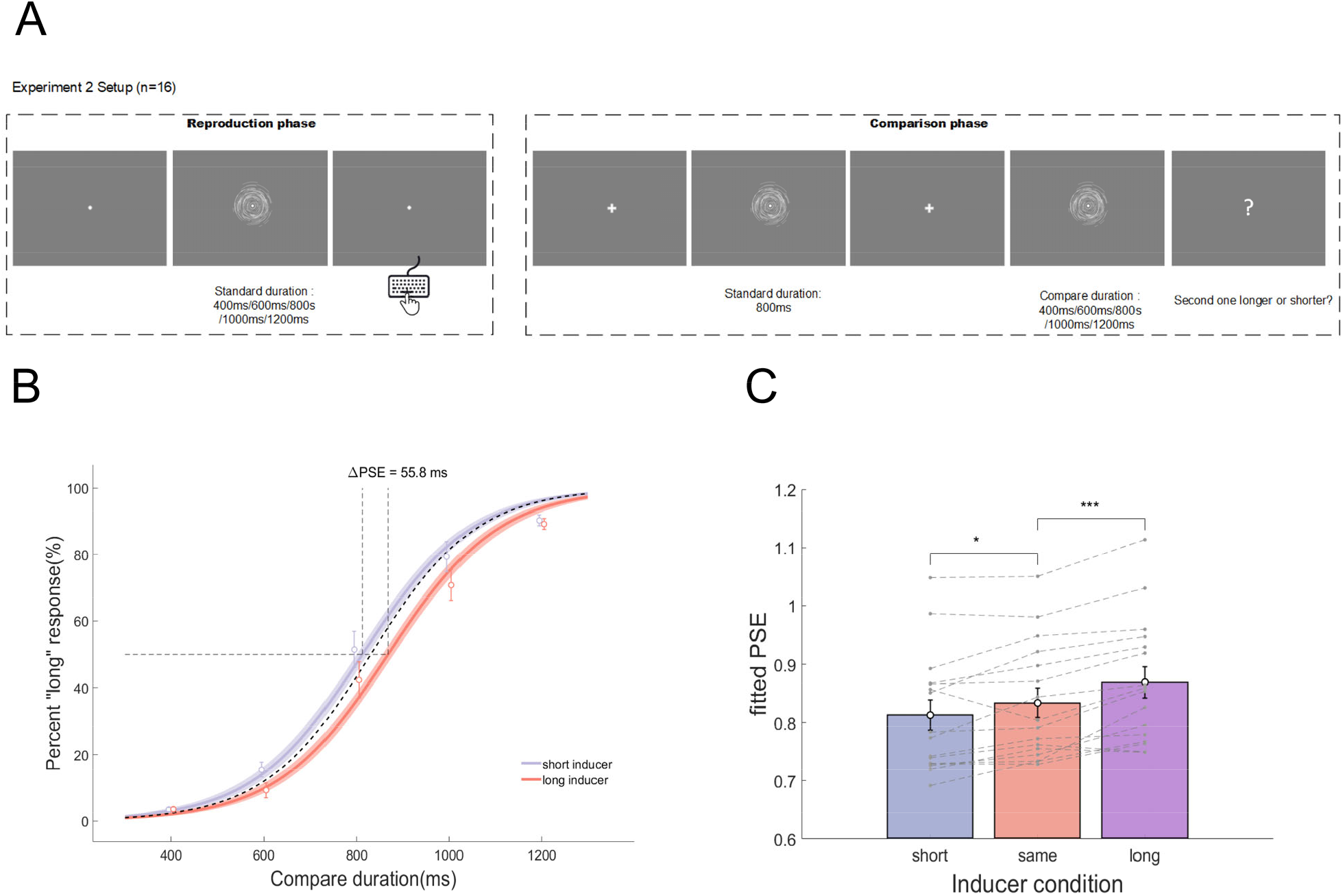
(A) Schematic procedure of experiment 2. Each trial consisted of a reproduction task and a comparison task. The reproduction task started with a circle central fixation point. A ripple pattern was then presented on the screen for a certain sample duration. Participants were instructed to press and hold a key for a period that matching the standard duration. Then the fixation point changed into a cross, indicating the following phase turned into a comparison task. Two ripple patterns were sequentially presented on the screen, with the first one as the standard duration and the second one as the compare duration. After the second ripple pattern disappeared, the central fixation turned into a question mark, promoting the participant to judge is the second one longer or shorter. No feedback was given on either of those two tasks. (B)Group level psychometric curve of different inducer condition. Different colors represent different inducer conditions, with purple denotes the short inducer condition (400 ms and 600 ms), red denotes the long inducer condition (1000 ms and 1200 ms). The dash line represents the equal inducer condition (800 ms). Shaded area represents 95% confidence interval band separately for each inducer condition. Hollowed circle represents the average proportion of “long” response under corresponding compare duration across all subjects. Error bar represents standard error. (C) Subject level PSEs of different inducer condition. Different colors represent different condition. The filled dots represent individual data. Error bar stands for standard error.

Before the formal experiment, participants completed a practice session with 20 trials in order to familiarize themselves to the task. Participants completed 8 separate experimental blocks in total, each of which contained 100 trials. After participants completed the 8 experimental blocks, they would be given another additional block. In the additional block, they only need to perform the comparison task without the reproduction phase. We use this block as a baseline. There were 100 trials in the additional block. All stimuli were identical with the two-phase task. All but one subjects completed this additional block.

After participants completed all experiment blocks, they were asked to rate the difficulty of the comparison task in the first 8 blocks and in the last block on a 7-point Likert scale.

### Analysis

In order to analyze the biases during the comparison phase, we fit psychometric curves to the behavioral data. The function we used was as follows:

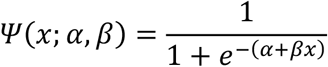

where *Ψ* denotes the proportion of subjects choosing a “longer” response, *α* denotes the position parameter, *β* denotes the slope of the curve, and *x* represents the difference between the standard and comparison stimuli. *α* determines the location of subjective point of equivalence (PSE), which in our hypothesis will shift if the reproduction task has a sequential effect on the following comparison task.

The raw data were split into two conditions according to the reproduction duration on the first phase. If the standard duration in the reproduction phase had a duration of 400 or 600 ms, that is, was shorter than the standard duration, then the trial was classified as “short inducer”, and if the standard duration in the reproduction phase had a duration of 1000 or 1200 ms, that is, was longer than the standard duration, then the trial was classified as “long inducer”. Trials on which the previous reproduction duration was 800 ms, that is, identical to the standard duration, were discarded from this analysis. In our hypothesis, the long inducer condition will shift the subject’s PSE rightward, and the short inducer condition will shift the subject’s PSE leftward. The subtraction of PSEs on the two conditions *ΔPSE* = *PSE*_*long*_ - *PSE*_*short*_ was classified as the bias induced by the reproduction, with positive values representing the attraction effect and negative values representing the repulsion effect. For the analysis of different inducer durations, we use the same approach.

On group level, permutation test was adopted to test whether the *ΔPSE* was significantly higher than zero under different conditions. Taking the short inducer and long inducer conditions as an example, we first shuffled the condition labels of the responses and then fit it with the psychometric function to calculate the *ΔPSE*. This process was repeated 10,000 times to obtain the distribution of *ΔPSE* under the null hypothesis, and the p-value was the proportion of *ΔPSE* under the null hypothesis that was greater than or equal to *ΔPSE* under the empirical distribution. The analysis for different inducer conditions and n-back analysis were performed in the same way as above.

We also perform the fitting process on the individual level and obtained subject-wise PSEs on different inducer condition. We then perform repeated-ANOVA to test whether the duration of inducer would cause a PSE shift. All tests were two-sided with a significance level *α* = 0.05 and Tukey correction was used for multiple comparisons.

## Results

In experiment 2, a reproduction task and a comparison task were interleaved to exclude the possibility of motor hysteresis. Through manipulating the immediate prior experience, we are expecting to observe a PSE shift for different conditions. As shown in Figure 2.B, we plot the proportion of “long” response against the comparison interval, and we fit the psychometric curves on the group level for different inducer conditions.

Given a previous shorter reproduction duration, the PSE of comparison task shifted to the left, and if preceded by a longer reproduction duration, the PSE shifted to the right. On group level, the *ΔPSE* reached 55.8 ms between the two conditions. Permutation test showed that the *ΔPSE* between the long and short conditions was significant (*p* < .001), so did the PSE shift between long and equal condition (*p* < .001), and PSE shift between short induction condition and equal condition (*p* = .0464). These results were shown in Figure 2.B.

We also conducted the analysis on the subject level. First, we fitted the psychometric curve for each individual subject and obtained their PSEs on different inducer conditions. Then we performed r-ANOVA and the result was shown in Figure 2.C. PSEs among different inducer conditions are different(F_(2,30)_=27.5, *p* < .001), PSEs of inducer with same length were longer than those of shorter inducer(t_(15)_=2.68, *p* = 0.043), while PSEs of inducer with same length were shorter than those of longer inducer(t_(15)_=5.13, *p* < .001).

Next, we want to ask if the PSE shift could be greater if the difference between the reproduction interval and standard interval in comparison is larger. And the result showed that a reproduction duration of 400 ms would cause the PSE shift 5 ms more left than a duration of 600ms (*p* = .333), and a duration of 1200 ms would cause the PSE shift 8 ms more rightward than a duration of 1000 ms (*p* = .273), but these results did not reach significance.

We also evaluated the serial dependence of reproduction tasks between trials. Using the same protocol as in experiment 1, we examined whether the reproduction task in the last trial could have an effect on the reproduction task on the current trial. The result showed no such inter-trial serial dependence exist (*p* = 0.11).

### Experiment 3

In the third experiment, we further investigated how a delay period would influence a potential bias. To this end, we manipulated the time interval between the stimuli presented and the subject’s response. We expected that the amplitude of serial dependence differs at different delays, and that the effect would be likely exhibit an increasing strength with longer retention time of information in working memory.

## Materials and Methods

### Participants

Sixteen healthy volunteers (8 males and 8 females, 18-30 years old, mean age = 22.40 ± 2.41) with normal or corrected to normal vision participated in Experiment 1. All subjects reported not taking any psychotropic drugs before the experiment. Before the experiment the participants have given written informed consent; this experiment and experiments 2 and 3 were approved by the institutional review board at Peking University. Subjects were screened for red-green color blindness before conducting the experiment. Financial compensation was given upon the completion of the experiment. All subjects finished four blocks of experiments separated by two days.

### Apparatus

The visual stimuli were presented on a 27-inch monitor equipped with a GeForce 1080 graphics card. The computer was equipped with Windows 10 operating system, with a refresh rate of 100 Hz, and a resolution of 1920 × 1080 pixels. Participants were seated 70 cm from the monitor in a dark soundproof room, participants’ heads were stabilized using a chin rest. The brightness of the screen was controlled in an appropriate range and kept constant, and a Dell keyboard is used for response.

### Stimuli and procedure

The experimental procedures were generated using MATLABR2021b and PsychToolBox 3.0.14 toolkit (Mathworks, Natick, MA; Brainard, 1997; Pelli, 1997). A scheme of the task is shown in Figure 3.A. At the beginning of each trial, a gray center fixation with 0.5 degrees in diameter was presented on a black screen. After a period of 0.75 s to 1.25 s a pair of brief fan-shape stimuli (see below for stimulus descriptions) were presented sequentially on the screen. Subjects were instructed to reproduce the intervals between the pair of stimuli with their dominant hands. The standard interval ranged from 540 ms to 1260 ms in ten arithmetical equal steps. After the second stimulus disappears, a delay period (0s, 1s, 3s, and 6s) was introduced before the subject’s response. After a certain delay period, the fixation circle transformed into a fixation cross to prompt the subject to respond as soon as possible. Subjects responded by hold the spacebar for a duration that matches the standard interval. Adaptive feedback was given by the color of fixation at the end of each trial. Reproduction accuracy was calculated by 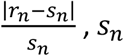 presents the standard duration of the *n*th trial and *r_n_* represents the reproduced duration of the *n*th trial. The initial threshold is set at 30% and will be updated adaptively with the subject’s performance. Trial with a reproduction accuracy below the threshold was given “correct” feedback (green fixation point), otherwise a red fixation point was given as “incorrect” feedback. The feedback signal lasts for 100ms. This feedback ratio was controlled using a one-up, one-down staircase method that adds or subtracts 3 for each incorrect or correct trial, respectively. After the feedback, the fixation changed into a gray circle and indicated the beginning of the next trial. Participants completed 4 separate experimental blocks, each of which contained only one condition of delay time. Before each block, participants completed a practice session with 20 trials in order to familiarize themselves to the task. After the practice session, participants completed a total of 200 trials in each block. Each standard interval was repeated for 20 times. Each block was divided into 4 sessions and there was a resting period between adjacent sessions. The presentation order of intervals within a session was randomized. The order of delay period across participants was counter-balanced using a Latin square method. Participants completed all 4 blocks on two separate experimental days.

**Figure 3.**
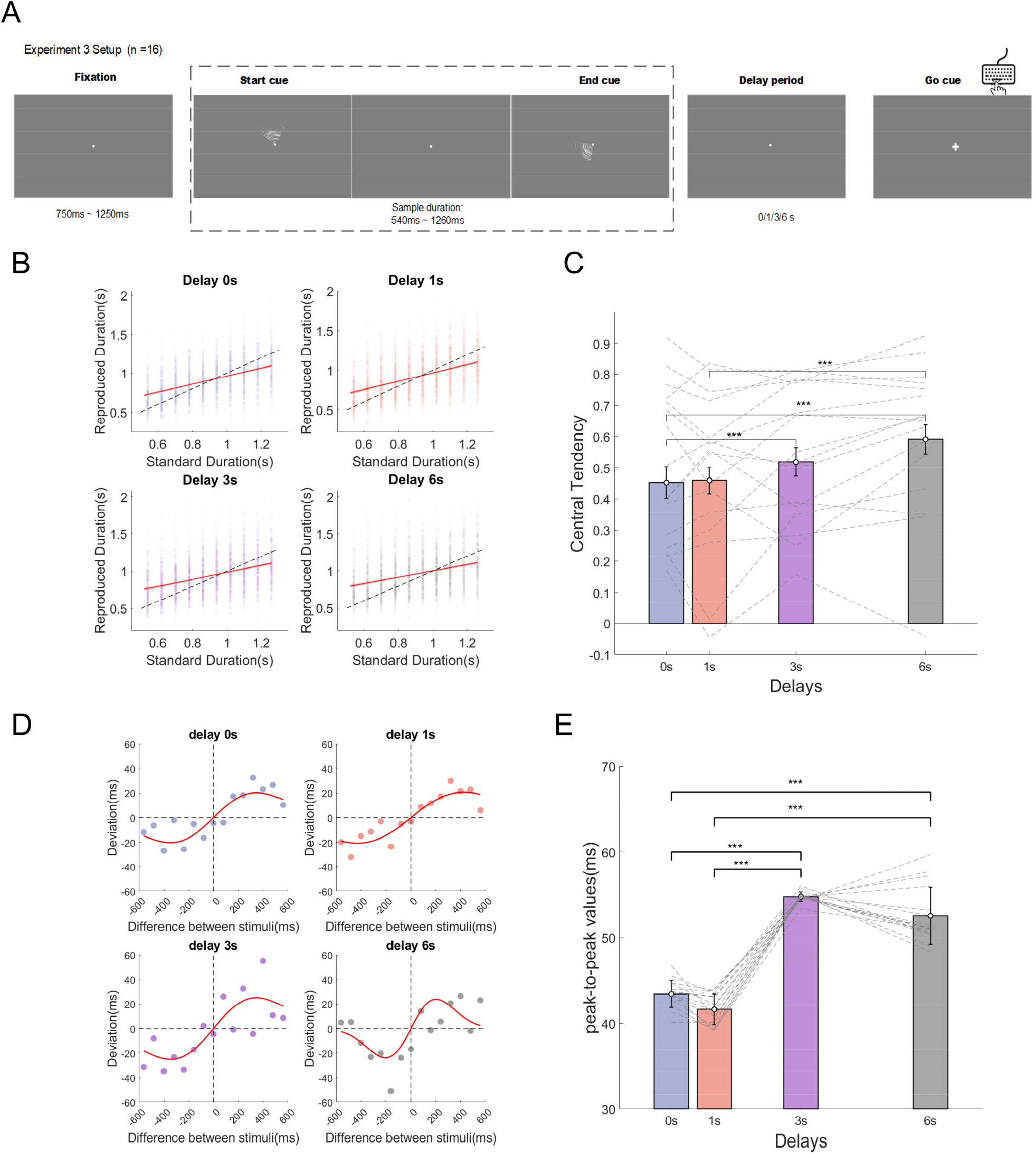
(A). Schematic procedure of experiment 3. Two brief flashes were sequentially presented and participants made button presses after a varying waiting time. Correct or incorrect feedback was presented after the response on each trial. Each trial started with a fixation point in the center the screen. After a random fixation time, two fan-shape patterns will flash on the screen for 100ms subsequently, the standard duration is indicated by the interval between the two patterns. After the presentation of the second fan-shape, there was a waiting time. Subjects were instructed to withhold their response until they saw the fixation point turned into a cross. The task was to replicate the standard duration by pressing a key. A feedback signal was given on each trial.(B) Central tendency for different delays. Each dot represents one subject. Red solid line represents linear regression fit to the data. Dashed line represents the reference line of y = x. (C) The strength of central tendency under different delays. Different colored bars represent the average values under corresponding condition. Each dashed line represents one subject. Error bar represent standard error. (D)Fitted DoG curve for different delays. Colored dot represents the group mean under corresponding condition. The red curve represents the fitted DoG curve between deviation(the reproduced duration minus the mean reproduced duration under the same condition) and difference between successive standard stimuli(the standard duration from previous trial minus the standard duration from current trial).(E) The strength(peak to peak values) of the best-fitted DoGs under different delay conditions. Each bar represents the group average under different delay conditions. Error-bar represents standard error.

Previous studies have found that successive presentations of the identical visual stimulus would cause repetition inhibition, resulting the following duration to be perceived as shorter (Matthews, 2011). Therefore, in the present experiment, we pre-generated 20 different ripple-like stimuli and present in each trial a different pattern. The stimulus luminance was kept consistent across trials. In Experiment 3, we used a 1/4 sector of the ripple stimuli to indicate the beginning and end of the standard interval. The two sectors were controlled to not overlap in space.

### Analysis

We removed the first trial in each session since there was no preceding trial. Trials with reproduced durations shorter than 0.3 s or longer than 1.5s or deviated more than 3 SDs from each condition’s mean were seen as outliers and excluded from all analyses (126 trials in total, 9.8 ‰). Outliers or unattended trials usually drastically affected the results, since subjects underwent an attention shift and the current stimulus did not enter the perceptual process or was processed incorrectly. In previous studies, it was found that only the correct response trials had an effect on the perceptual bias of the current trial, while the incorrect trials had no effect on the perceptual bias (Braun et al., 2018). Thus, pre-exclude abnormal trials were necessary for further analysis.

For the analysis of central tendency, the reproduced intervals were linearly regressed to the standard intervals, and we used one minus the linear regression coefficients as indices of the central tendency. The range of central tendency was then between 0 and 1, with larger values representing larger central tendency.

As for the analysis of serial dependence, there are two different types of metrics: absolute serial dependence (ASD) and relative serial dependence (RSD)(Glasauer & Shi, 2021; Fischer & Whitney, 2014). ASD is calculated based on the response error of the current trial and the standard stimulus of the previous trial, while RSD uses the response error of the current trial and the standard stimulus difference between of the previous trial and the current trial (Alais et al., 2017; Cicchini et al., 2017). In the following, we will use RSD as an indicator of serial dependence, as RSD has a more realistic reflection for nonlinear characteristics of our experimental data and has better mathematical intuition. We also performed ASD analysis in the later part as for a comparison purpose with a previous study. It should be mentioned that those different metrics didn’t differ in the conclusion. At the same time, to prevent false positive results, we also performed sanity check to ensure the correctness of the analysis method (see the supplementary for more information). The calculation process for serial dependence is same as in experiment 1.

## Results

To estimate the central tendency, we calculate the difference between one and the regression coefficient of reproduction duration as a function of the standard duration for different delays conditions. The central tendency effect enhanced significantly as the delay increased (*F*_(3,45)_ = 12.4, *p* < .001, *η*^2^ = 0.087), (Figure 3.B & C). The mean central tendency for delay 0 s was 0.452 ± 0.205, delay 1 s was 0.459 ± 0.171, delay 3 s was 0.501 ± 0.184, delay 6 s was 0.591 ± 0.189 (mean ± SD), and all results were significantly different from zero (p<.001). Post-hoc comparison showed that the central tendency was significantly different between delay 0 s and delay 3 s(*t*_(l5)_ = -3.383, *p* = 0.025), delay 0 s and delay 6 s(*t*_(l5)_ = -4.393, *p* = 0.003), delay 1 s and delay 6 s(*t*_(l5)_ = -5.213, *p* < .001). All other comparisons were not significant. The result suggested that central tendency was modulated by the delay time, with there seemed to be two clusters for the four delays. There is no difference between delay 0 s and delay 1 s, no difference between delay 3 s and delay 6 s. But the two clusters differed significantly.

For the serial dependence effect, we fit the deviation as a function of the difference between the standard stimulus using the DoG model. The analysis was performed on group level. The model fitting was conducted separately for different delays and the result is shown in Figure 3.D. We characterized the strength of serial dependence using the peak-to-peak value, defined as the difference between the highest and lowest values of the DoG curve. To estimate the variations of peak-to-peak values at delay conditions, we used the JackKnife resampling method, where each participant was left out once from the collapsed group data. We then compare these estimations using one-way ANOVA.

Similar to the central tendency effect, one-way ANOVA showed that the magnitude of the serial dependence was significantly increased as the delay increased (Figure 3.E). Games-Howell Post-hoc tests for different delays showed that, the p2p values was significantly different between delay 0 s and delay 3 s(*t*_(l8.3)_ = -27.1, *p* < 0.001), delay 0 s and delay 6 s(*t*_(2l.5)_ = -9.88, *p* < 0.001), delay 1 s and delay 3 s(*t*_(l7.6)_ = -28.0, *p* < 0.001), delay 1 s and delay 6 s(*t*_(23.l)_ = -11.54, *p* < 0.001). There was no statistically significant difference in p2p values between delay 0 s and delay 1 s (*t*_(29.9)_ = -2.36, *p* = 0.108) or between delay 3 s and delay 6 s (*t*_(l5.8)_ = 2.66, *p* = 0.073). This result suggested that the serial dependence started to rise shortly after the offset of stimuli and saturate in a later period.

## The relationship between central tendency and serial dependence

Another famous context influence in temporal reproduction task is the Vierordt effect, i.e., overestimation of relatively short and underestimation of relatively long temporal intervals (e.g. Pöppel, 1971/72, 1997; Vierordt, 1868). This effect is also assumed to reflect central tendency. Several previous models proposed that central tendency and serial dependence might result from the same underlying mechanism (Petzschner & Glasauer, 2011; Wiener et al., 2014). Petzschner and Glasauer developed an iterative Bayesian model that updates the internal reference, the prior, on a trial by trial basis and argued that the Vierordt effect is a by-product of an optimal strategy that aiming to improve the reliability by utilizing extra cues. Though these two phenomena both fit in the Bayesian framework, the underlying assumptions are distinct. The Vierordt effect, or regression to mean, considers entire distribution from which the stimuli are drawn. Serial dependence takes place in a real-time trial-by-trial manner, and the effect size is comparably smaller than Vierordt effect. In other words, the Vierordt effect assumes a static prior throughout the experiment, and the successive stimuli are independent of each other. On the contrary, an iterative process is essential for the generation of serial dependence, the consecutive stimuli are believed to have temporal continuity. In a recent study by Glasauer and Shih (2021) the authors proposed a two-stage model that combined the static model with a simple iterative model. The model assumes that the mean of the stimulus distribution in each trial varies in a random walk manner. This imparts the auto-correlation feature to the model and a random Gaussian noise with fixed variance is superposed on that. They also derived an upper and lower boundary for the relationship between central tendency and serial dependence. The lower boundary was confined by a static model, which predicts that regardless of the central tendency, the serial dependence will stay at zero. The upper boundary is on the other extreme and predicts that the serial dependence has a deterministic relationship with the central tendency.

The relation of serial dependence and central tendency for all individual participants is depicted in Figure 4. According to the two-state model, all points shall lie within the shaded area. What we observe here was that there were two obvious outliers that showed relatively large serial dependence while kept a small central tendency. Though most of the participants were within the range, the-two stage model partially failed to explain the two outliers we observed. Alternatively, we computed the first-order correlation between the ASD and central tendency by fitting a regression line, the result showed there was significant negative correlation between central tendency and serial dependence (regression coefficient = -0.2558, *p* = 0.0026, *AdjustedR*^2^ = 0.4517). This result might imply that there is a balance between the two strategies that subjects use when encounter with a duration reproduction task. On one extreme are subjects with low serial dependence but high central tendency, their hypothesized model about the world assumes that the stimuli was drawn from a fixed distribution with a fixed mean and no relationship between successive stimuli. On the other end are subjects who implicitly believe that there is temporal continuity to some extent, i.e., that the world is changing slowly; it is an optimal strategy to just integrate over short period than assume a static prior. Most of the subjects showed a combination of the two strategies though we could observe some extreme cases in the figure. Furthermore, the results that both serial dependence and central tendency were modulated by the delay time suggests that there might be an indirect connection between those two effects through some mediating mechanism.

**Figure 4.**
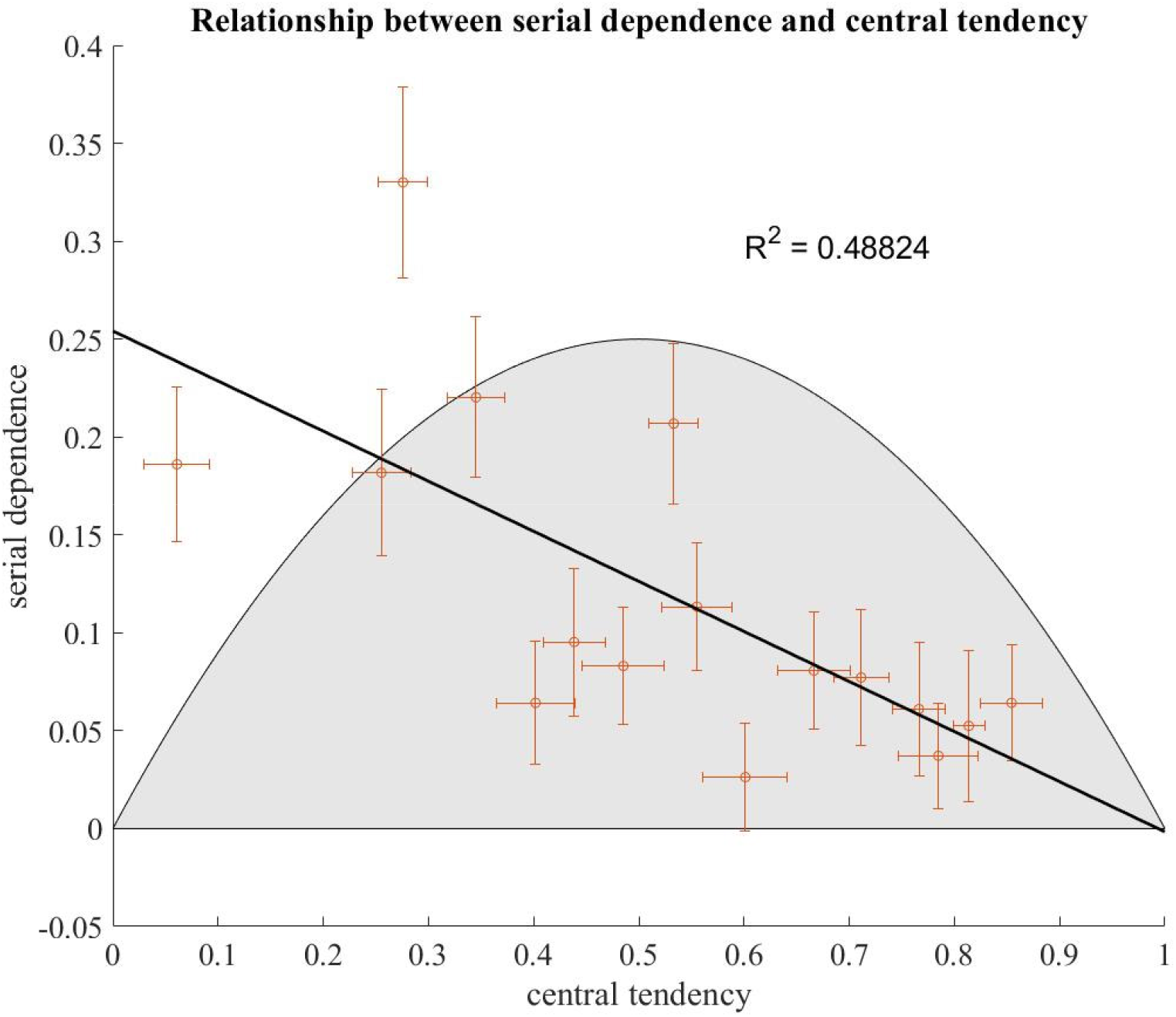
The relationship between central tendency and absolute serial dependence. Each open dot represents an individual subject. The shaded area are predicted plausible regions of the two-stage model from (Glasauer and Shi 2021a). Error bar represents standard error. The solid black curve represents the first-order linear regression line.

### General Discussion

Our prior experience can affect how we perceive subsequent events. Previous studies proposed that the serial dependence might be a common processing mechanism and includes various cognitive functions. It is theoretically worthwhile to study the serial dependence in temporal perception and how different functions are potentially involved. To this end we conducted a series of experiments to examine serial dependence in temporal perception. We hypothesize that for temporal perception, subjects do not only rely on real time perception of stimuli, but the information about the context is also recruited to create a prior for the current temporal processing.

In the present study, we addressed these questions by investigating the serial dependence using temporal reproduction and temporal comparison tasks. We showed that the prior experience influences both the following reproduction task and the comparison task. In both these tasks, the bias was attracted towards the preceding stimulus, i.e., a positive serial dependence. In experiment 1 we extended previous research, and found that perception without a motor response could also cause although a weaker serial dependence. In experiment 2 we demonstrated that the bias did not rise from motor hysteresis, i.e., the tendency to repeat the motor response in the last trial; instead, it altered the perception of the interval in a task that does not include a motor response. And in experiment 3, we demonstrated that this effect occurred regardless of the delay period, and the magnitude of the effect increases with delay time.

### Perception without motor response showed a weak attractive effect

Previous studies on serial dependence showed inconsistent results when testing serial dependence in perception without motor response in visual perceptions (Manassi et al., 2018; Pascucci et al., 2019). It is even more difficult to predict this result under the context of the temporal perception since previous results on duration judgment implied opposite sequential effects. When participants are continuously exposed to one duration and suddenly perceive a new one, they tend to overestimate the difference between the old and the new durations, known as the adaptation effect, and predicting negative serial dependence (Heron et al., 2012; Li et al., 2015, 2015, 2017; Shima et al., 2016; Walker et al., 1981). However, studies in the duration judgment tasks show that the estimation of the current comparison duration is attracted by the one in the previous trial (Lapid et al., 2008; Rammsayer & Ulrich, 2012). Moreover, inconsistent results were found in the recent studies regarding whether the passive observations have any influence on regression-to-mean of the duration reproduction in different studies (Roach et al., 2017; Zimmermann & Cicchini, 2020), which make it more confusing whether motor response is necessary to construct the internal reference frame. In Experiment 1, we indeed observed a significant attraction from the previous stimuli duration instead of an adaptation effect after amplifying the effect of perception by repeatedly presenting durations either all longer or all shorter than the target sample duration. This attraction effect confirms that the perception itself is sufficient to influence the internal reference. However, the impact of perception is much weaker than the reproduction and can only be detected when repeated five times, which perhaps explained no impact of passive observation on central tendency in some previous study.

### Serial dependence as a common mechanism in different paradigms

In experiment 2, we used a two-stage task to show that the serial dependence was not specifically linked to the temporal reproduction task. Methods to evaluate temporal perception are extensive, and a central problem that lies among those massive methods is the question whether they reflect a common factor of temporal perception or whether the measurements are task-specific (Bao et al., 2015; Pöppel, 1971/1972; Zhou et al. 2014). Here we used two classical temporal duration paradigms and found that positive serial dependence existed regardless of different paradigms. Furthermore, the strength of serial dependence on a group level was also comparable with experiment 3. In experiment 3, the peak-to-peak values for delay 3 s was 54.8 ms, for delay 6 s 52.5 ms, while the PSE shift here was 55.8 ms. The two different tasks converging to a similar serial dependence strength suggest that there might be a central mechanism participating in the integration of temporal adjacent events.

It should be noted that in the temporal comparison task there was a commonly observed bias, i.e., a time-order error. With brief intervals the first interval is overestimated and/or the second one is underestimated. According to previous research, the comparison is rather accurate if two stimuli are presented for a temporal interval of two or three seconds (Pöppel, 1971/72, 1997; von Steinbüchel et al., 1996). Since we adopted a very short inter-stimulus interval in the comparison task, we exclude the possibility that the bias we observed here was contaminated or caused by time order error.

Several previous studies suggested that the generation of the context bias was related to the task itself. In a temporal reproduction task, the effect could be detected only when the previous stimulus was accompanied with a motor response. In a temporal bisection task, no motor response was required, the effect emerged automatically without any further control (Cai et al., 2017; Wehrman et al., 2018). This indicates that the task demand might play a role in the generation of serial dependence. It is interesting to ask whether the context bias generated by one task will affect another different task in the following, or whether the effect can only carry over within the same kind of tasks. According to our results, the former holds true. Taken together, in experiment 2 we demonstrated that serial dependence can also be transferred between different temporal tasks. This adds general support for the robustness of the effect and the evidence accumulation process between different temporal proximity tasks provides a new perspective on the mechanisms of serial dependence.

### The dependency of serial dependence on delay time

By manipulating the delay time between the stimulus and the response, we could assess the temporal course of trial history. We observed an increasing magnitude of serial dependence with lengthened delay time, similar to previous results in visual tasks (Bliss et al., 2017; Fritsche et al., 2017). Our findings suggest an increasing magnitude of serial dependence over longer delay periods, which may reflect a relation with working memory processes. If the serial dependence merely would happen at a perceptual level, once the perception is formed, it should presumably not change with increasing delay time. The alternative hypothesis is that if the serial dependence has a mnemonic component, it should scale with an increasing delay period. Our results support the latter statement. Although the effect we observed in experiment 3 saturated after 3 s and did not continue increasing with delay time, this does not contradict previous findings that the effect saturates at around 6 s (Bliss et al., 2017). This was possibly due to the different trial length of the tasks. In a temporal reproduction task, the average trial length is longer than a visual task. Taking this into consideration, the interval between the offset of response in the last trial and the onset of response in the current trial would be approximately same. The statement that serial dependence in temporal perception involves a mnemonic component does not exclude the possibility that it could also has a perceptual component. In fact, in experiment 1, we observed a positive serial dependence after several no-action trials.

The involvement of working memory in the creation of serial dependence is supported also from research on time windows. Experiment 3 showed that two clusters up to 3 seconds and beyond 3 seconds. Ample evidence has been collected about a pre-semantic time window of approximately 3 seconds that underlies cognitive processes as a logistic function (Bao et al., 2015; Pöppel, 1997, 2009; Zhou et al. 2014). Using the reproduction paradigm (Mu et al., 2022) behavioral experiments with healthy adult subjects (Pöppel 1971/72) or children (Szelag et al., 2002), with brain injured patients (Kagerer et al., 2002; von Steinbüchel et al., 1996) or autistic children (Szelag et al., 2004), and electrophysiological studies (Chen et al., 2015; Elbert et al., 1991; Wang et al., 2016) have substantiated the functionality of a time window of approximately 3 seconds. Thus, the data obtained in experiment 3 support on the one hand the notion of this time window and on the other hand furthermore provide an explanation of the two clusters in serial dependence as one is dealing with two different neural processes.

## Conclusion

One crucial question about serial dependence was what role the perception component plays in it. By accumulating no-action trials, we managed to prove that perception without motor response was sufficient to generate serial dependence but with a relatively weaker attraction. We further tested whether serial dependence could bias the perception in a different task following a reproduction task. The results confirmed the robustness of the effect and demonstrated that it could be transferred between different tasks. Further, we used the delayed temporal reproduction paradigm and found that for brief presentation of interval, only attractive serial dependence emerged with the absence of repulsive adaptation. By checking the time course and strength of serial dependence, we argue that the serial dependence involves the participation of working memory. Our study showed that the serial dependence might not be a goal-directed behavior merely, but rather a shared mechanism across different tasks.

## Acknowledgements

This work was supported by National Natural Science Foundation of China (Projects 31771213 and 31371018) to Yan Bao.

## Author contributions

S.C. and T.W. designed the experiment. S.C. and T.W. collected the data. S.C. and T.W. analyzed the data. S.C. prepared the figures. S.C., T.W., and Y.B. involved in writing the manuscript with the initial draft written by S.C.

## Competing interests

The authors declare no competing interests.

## Additional information

The experiment code and analysis code from this study are both freely available online at Github

## Notes

### Competing Interest Statement

The authors have declared no competing interest.

